# Multimodal immobilization of second-instar *Drosophila melanogaster* larvae using PF-127 hydrogel and diethyl ether for calcium imaging

**DOI:** 10.64898/2026.03.19.713048

**Authors:** David Reynolds, Eliana Artenyan, Hayk Nazaryan, Edward Shanakian, Elva Chen, Vaheh Abramian, Ava Ghashghaei, Kian Sahabi, Fuad Safieh, Nicolas Momjian, John Sunthorncharoenwong, Javier Carmona, Katsushi Arisaka

## Abstract

Motion artifacts remain a barrier to in vivo calcium imaging in *Drosophila melanogaster* larvae. Here, we evaluate a multimodal immobilization approach that combines a Pluronic F-127 (PF-127) hydrogel with brief diethyl ether vapor exposure (5 minutes, 25°C) and compare it against hydrogel-only immobilization using custom MATLAB-based analysis software that performs NoRMCorre rigid motion correction.

In wide-field GFP recordings at 1 Hz over approximately 60 minutes (*N* = 15 per group), the multimodal condition significantly reduced motion across all three core metrics after FDR correction (all *q <* 0.001), with large effect sizes for mean speed (Hedges’ *g* = −1.18) and median step size (*g* = −1.36). In a secondary analysis of the first 30 minutes, uniformly large effect sizes (|*g*| = 1.10–1.51) were observed, consistent with stronger initial chemical immobilization that partially wanes over the recording period.

We implemented a dual-flag quality control system that distinguishes motion data reliability from ROI detection eligibility. Control calcium recordings (33.33 Hz, ∼5 minutes; *N* = 23) yielded 368 ROIs with a mean SNR 30.4 ± 16.9 and an event rate of 0.228 ± 0.113 Hz. Experimental recordings (*N* = 21) yielded 295 ROIs with SNR 18.0 ± 10.6 and event rate 0.309 ± 0.188 Hz. SNR was higher in controls (Cliff’s *δ* = 0.50, *p <* 0.001), while event rate was modestly higher in the experimental group at the ROI level (*δ* = −0.22, *p <* 0.001), though this difference did not reach significance at the sample level, suggesting altered but not suppressed calcium dynamics. These results support a practical, accessible immobilization workflow for larval calcium imaging.

**Highlights:**

- Brief ether + hydrogel approach reduces larval motion 85–91% vs. hydrogel alone
- Dual-flag QC system separates motion reliability from calcium ROI eligibility
- Calcium event rates not suppressed under multimodal immobilization
- Complete MATLAB pipeline for motion analysis and calcium imaging provided
- Accessible protocol requires only standard laboratory supplies

## 1 Introduction

### 1.1 *Drosophila* larvae as a model for neural imaging

*Drosophila melanogaster* has served as a foundational model organism in genetics and neuroscience for over a century, offering precise genetic manipulation, extensively characterized biology, and experimental accessibility [Bellen et al., 2010]. The larval stage is particularly well-suited for in vivo neural imaging: its nervous system is compact, with approximately 3,016 neurons in the brain recently mapped at full synaptic resolution [Winding et al., 2023], yet capable of supporting complex behaviors. The semi-transparent cuticle permits optical access without invasive surgery, enabling experiments that directly relate circuit architecture to neural activity.

The GAL4/UAS binary expression system [Brand and Perrimon, 1993] allows genes to be expressed in defined neuronal populations, and thousands of driver lines are available through stock centers such as the Bloomington Drosophila Stock Center. When combined with genetically encoded calcium indicators of the GCaMP family, whose jGCaMP8 variants exhibit half-rise times as fast as 2 ms and improved decay kinetics [Zhang et al., 2023], these tools enable optical monitoring of neural activity in genetically defined cell populations across the larval nervous system [Grienberger and Konnerth, 2012, Chen et al., 2013, Dana et al., 2019].

### 1.2 The motion artifact problem

Despite these strengths, in vivo calcium imaging in *Drosophila* larvae is often limited by residual movement during recording. Even in restrained preparations, peristaltic contractions, gut motion, and subtle tissue deformation introduce frame-to-frame shifts that compromise image registration and downstream analysis [Lemon et al., 2015, Heckscher et al., 2012]. Motion artifacts are particularly problematic for calcium imaging because fluorescence traces are extracted from fixed spatial regions. When neurons shift across pixels between frames, the measured signal may reflect spatial displacement rather than genuine changes in intracellular calcium. This can artificially alter signal amplitude, introduce spurious correlations between neurons, and reduce the reliability of downstream event detection. Small changes in the focal plane alter apparent brightness independent of calcium dynamics, further reducing signal-to-noise. Global movement can also introduce artificial correlations across the field of view, and larger shifts cause tissue to drift partially or fully out of frame.

Computational motion correction can reduce some of these effects. NoRMCorre [Pnevmatikakis and Giovannucci, 2017], which supports both rigid and piecewise rigid registration, aligns frames to a reference template and applies corrective shifts. However, registration cannot restore information lost when tissue moves out of the imaging plane and is limited in its ability to resolve complex tissue deformation. For these reasons, improving the physical stabilization of the preparation remains essential.

### 1.3 Existing immobilization approaches

Multiple immobilization strategies have been developed for larval imaging, each with distinct advantages and limitations. Microfluidic devices provide excellent mechanical stabilization. Ghaemi et al. [2017] designed microfluidic clamps that restrict CNS movement to approximately 10 µm over 350 seconds, Chaudhury et al. [2017] combined mechanical compression with mild cryo-anesthesia to achieve submicron stability, and Ghannad-Rezaie et al. [2012] developed chips for imaging neural injury responses using mechanical compression with CO_2_ delivery. However, these approaches require soft lithography fabrication, cleanroom access, and in some cases specialized cooling or gas delivery systems, limiting their availability to well-equipped laboratories.

Thermoreversible hydrogels offer a more accessible alternative. Pluronic F-127 (PF-127) is a triblock copolymer that exists as a low-viscosity liquid below ∼15°C and forms a transparent gel at higher temperatures, allowing larvae to be positioned in chilled solution and immobilized as it solidifies at room temperature [Dong et al., 2018]. PF-127 is bio-compatible, optically clear, reversible by cooling, and requires no specialized equipment. However, the mechanical restraint provided by hydrogel alone may be insufficient to fully suppress larval peristaltic movements, particularly over extended recording sessions.

Chemical anesthetics directly suppress neural activity and muscle contraction. Kakanj et al. [2020] demonstrated that brief diethyl ether exposure (3–4.5 minutes, depending on instar) can immobilize larvae for up to 8 hours, with high survival rates and no specialized equipment required. However, general anesthetics, including diethyl ether, act through multiple mechanisms: potentiation of inhibitory GABA_A_ receptors, inhibition of excitatory NMDA receptors, modulation of two-pore-domain potassium channels, and effects on voltage-gated sodium channels [Hemmings et al., 2005]. In adult *Drosophila*, ether sensitivity depends substantially on the genotype of the *para* sodium channel gene, suggesting that sodium channel modulation is a primary mechanism in this organism [Tanaka and Gamo, 2001], though notably larval sensitivity to ether appeared less dependent on *para* genotype. These broad effects on neural signaling raise concerns about the validity of calcium imaging data collected under deep anesthesia.

Cold anesthesia avoids chemical perturbation but introduces its own confounds: synaptic transmission, channel kinetics, and circuit function are temperature-dependent, and imaging at reduced temperatures captures activity under non-physiological conditions [Tang et al., 2010].

### 1.4 Rationale for a multimodal approach

Given that each immobilization method presents trade-offs when used alone, we reasoned that combining approaches could achieve more effective immobilization while mitigating individual limitations. Specifically, we hypothesized that brief diethyl ether exposure could provide initial chemical suppression of movement, which would then be maintained by the mechanical restraint of a PF-127 hydrogel matrix as the anesthetic effects dissipate. This strategy aims to extend the effective imaging window while minimizing the total anesthetic burden.

We chose to compare this multimodal approach against hydrogel-only immobilization rather than against microfluidic or other technically demanding methods. PF-127 hydrogel is widely available, inexpensive, requires no specialized equipment, and represents the current accessible standard in many laboratories. Our goal was to determine whether adding brief ether pre-treatment to this established method could significantly improve immobilization quality as measured by quantitative motion metrics, and whether the resulting preparations remain suitable for calcium imaging.

### 1.5 Study objectives

The objectives of this study were to: (1) quantify motion in hydrogel-only and ether + hydrogel preparations using multiple motion metrics derived from frame-by-frame registration shifts; (2) develop transparent, automated quality control criteria that distinguish reliable motion tracking from tracking failures, and separately gate eligibility for calcium region of interest (ROI) extraction; (3) evaluate calcium imaging feasibility by comparing ROI detection yield, signal-to-noise ratio, and calcium transient detection between conditions; and (4) provide a documented, reproducible analysis pipeline supporting both motion quantification and calcium imaging processing.

## 2 Materials and Methods

### 2.1 Fly strains and genetic constructs

All experiments used *Drosophila melanogaster* larvae expressing genetically encoded fluorescent reporters under the GAL4/UAS binary expression system [Brand and Perrimon, 1993]. Pan-neuronal expression was achieved using the R57C10-GAL4 driver line [nSyb-GAL4; Pfeiffer et al., 2008]. Calcium imaging experiments used larvae expressing the genetically encoded calcium indicator jGCaMP8f [Zhang et al., 2023] pan-neuronally: R57C10-GAL4 crossed to ii20xUAS-IVS-RSET-jGCaMP8f (VK00005, 2nd generation insertion). Movement experiments used the same driver crossed to UAS-mCD8-GFP [y,w; UAS-mCD8-GFP; Lee and Luo, 1999], a membrane-targeted GFP reporter, for motion tracking without calcium-dependent signal fluctuations. The R57C10-GAL4 and UAS-jGCaMP8f lines were obtained from the Mark Frye laboratory (UCLA). The UAS-mCD8-GFP line was obtained from the David Krantz laboratory (UCLA).

### 2.2 Fly husbandry

Flies were maintained at 25°C with 40% relative humidity under a 12-hour light/dark cycle on standard cornmeal-molasses-agar medium prepared by the Drosophila Media Facility at UCLA. Pure stock lines were maintained in bottles with 20 females and 10 males. For experimental crosses, parent flies were renewed from the youngest available bottle of recently emerged adults every 7–10 days. Flies were anesthetized by cold exposure (4°C, 1 hour) for sex sorting. Crosses were established with 18 females and 9 males per vial, labeled with cross date, genotype, and initials of the person who set up the cross, and maintained at 25°C. Parent flies were transferred to fresh vials daily.

### 2.3 Larval staging and selection

Second-instar larvae were selected primarily by developmental timing. Larvae were collected from bottles 4–5 days after parent flies were introduced to fresh media; at 25°C, this timing yields predominantly second-instar larvae. As a secondary screen, CNS length was measured from wide-field fluorescence images using the straight-line tool in Fiji/ImageJ [Schindelin et al., 2012]. Images were calibrated at 0.260 µm*/*pixel (266 µm field of view at 1024 × 1024 resolution). CNS length was defined as the distance from the posterior tip of the ventral nerve cord to the dorsal margin of the central brain, excluding the optic lobes, measured along the longest apparent axis to account for variation in mounting angle. Larvae with CNS length clearly inconsistent with second instar, or with truncated, damaged, or ambiguous CNS boundaries, were excluded.

### 2.4 Immobilization protocols

Two immobilization conditions were compared. Both protocols used identical materials and handling steps except for the diethyl ether exposure.

#### Materials

Glass microscope slides (25 × 75 × 1.0 mm; Fisher Scientific), square coverslips (18×18 mm, thickness 0.13–0.17 mm; MUHWA Scientific), copper tape spacers (150 µm thickness; StewMac), Pluronic F-127 (Sigma-Aldrich) prepared as 25% w/v solution in cold, distilled water, HL3.1 saline [Feng et al., 2004], diethyl ether (Sigma-Aldrich), and a Coplin staining jar (Karter Scientific) used as the anesthetizing chamber.

#### PF-127 preparation

PF-127 powder was dissolved slowly in cold distilled water with continuous stirring to produce a 25% w/v solution. The solution was stored at 4°C, where it remains liquid. The solution gels upon warming above approximately 18°C.

#### Control condition (hydrogel only)

Second-instar larvae were extracted from media bottles using a paintbrush and transferred to a glass slide equipped with 150 µm copper tape spacers. Residual media was rinsed away with 0.5 µL HL3.1 saline [Feng et al., 2004]. 20 µL of pre-chilled 25% PF-127 solution was applied over each larva, a coverslip was placed on the copper tape spacers, which defined the compression height, and the slide was heated on a hot plate at 30–35°C for 3–4 minutes to accelerate gelation. Slides were then transferred to the microscope (Fig. 1).

**Figure 1:**
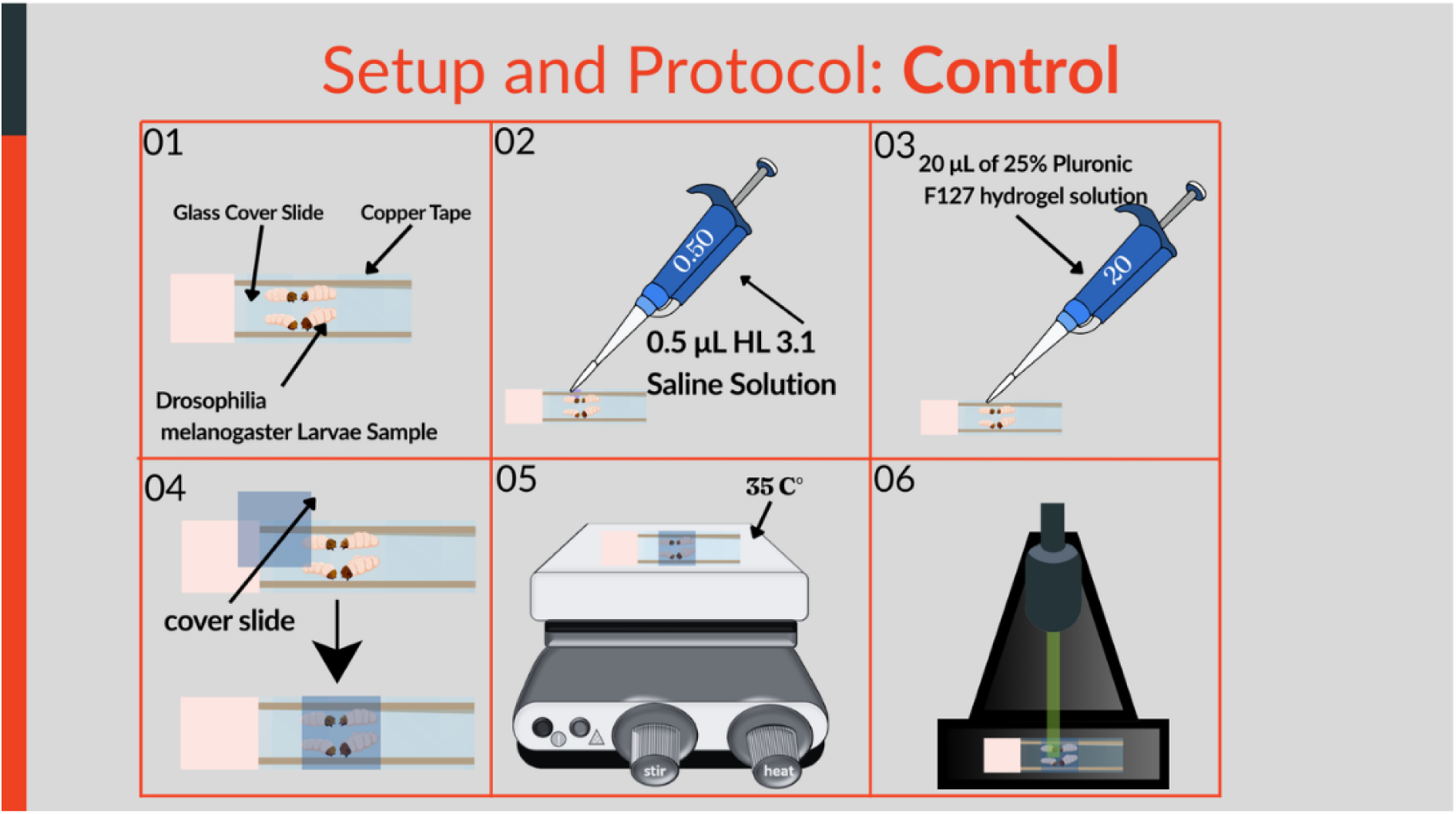
Control immobilization setup and protocol for imaging *Drosophila melanogaster* larvae. Larvae were placed on a glass slide bordered with copper tape and hydrated with 0.5 µL HL3.1 saline solution. A 20 µL drop of 25% Pluronic F-127 hydrogel was applied over the sample and covered with a glass coverslip. The preparation was heated to 30– 35°C to initiate hydrogel solidification, immobilizing the larvae for stable imaging under the microscope.

#### Experimental condition (diethyl ether + hydrogel)

Larvae were transferred to a wire mesh sheet, which was placed in a Coplin chamber containing diethyl ether (two cotton balls lightly moistened with approximately 5 mL total) for 5 minutes at room temperature (∼25°C). The mesh was then removed, and larvae were transferred to a glass slide. From the saline wash step onward, handling was identical to the control condition. Time from removal from the anesthesia chamber to imaging start was approximately 5–7 minutes (Fig. 2).

**Figure 2:**
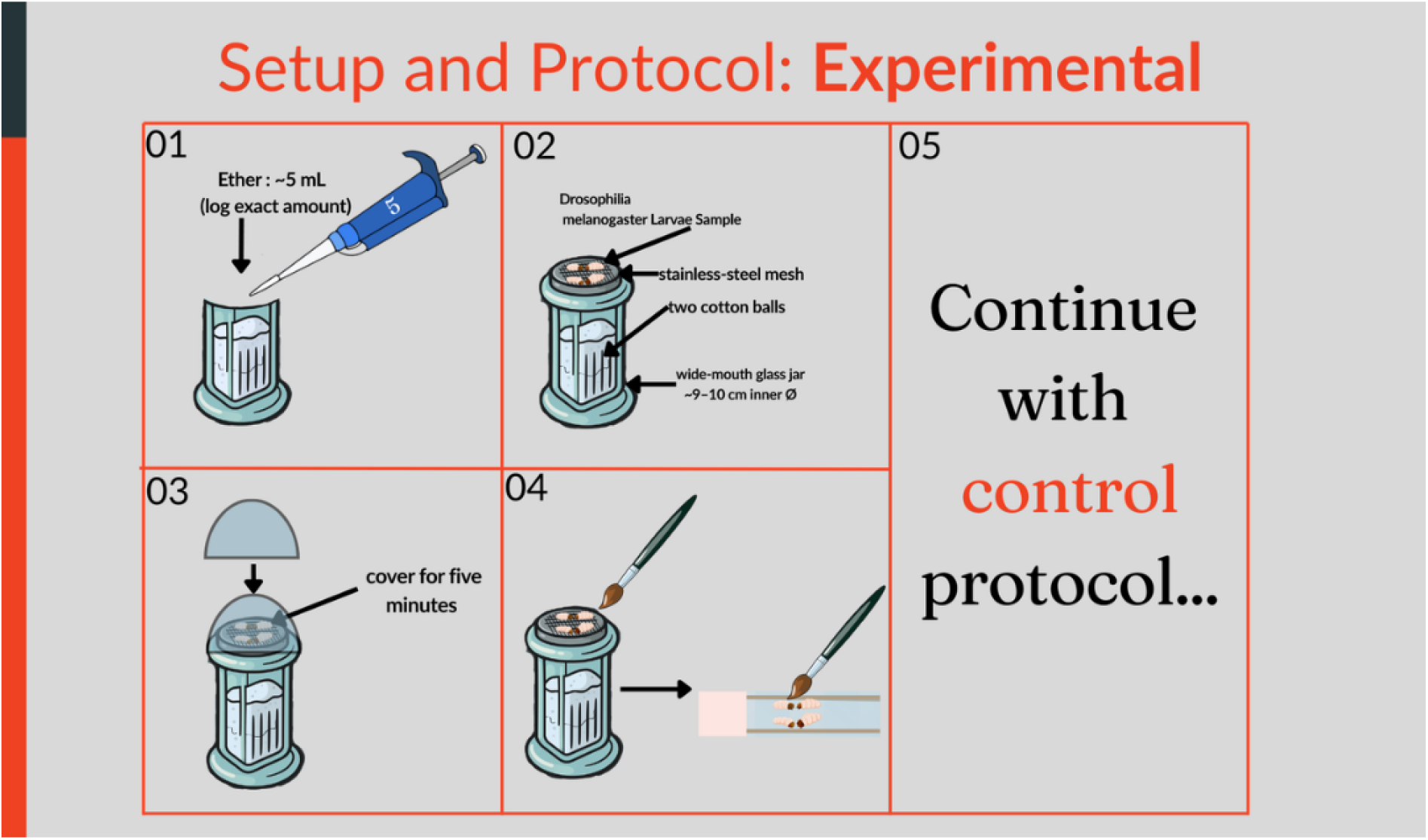
Experimental ether anesthesia protocol for *Drosophila melanogaster* larvae prior to immobilization and imaging. Approximately 5 mL of diethyl ether was added to a Coplin chamber containing cotton balls. Larvae were placed on a stainless-steel mesh above the ether source, and the chamber was covered for 5 minutes to induce anesthesia. After exposure, larvae were transferred to a glass slide, and the preparation was carried out using the control immobilization protocol (Fig. 1).

#### Safety precautions

Diethyl ether is an extremely flammable liquid and vapor (flash point −45°C) that can form explosive peroxides upon prolonged storage. All ether handling must be performed in a certified chemical fume hood, away from heat sources, sparks, and open flames. Protective gloves and eye protection must be worn during use. The Coplin chamber must be opened and closed within the fume hood. Ether stock must be stored in a flammable-safe cabinet, inspected for peroxide formation, and replaced regularly. Hands and exposed skin must be washed thoroughly after handling. Investigators should consult their institutional safety office before implementing this protocol.

### 2.5 Post-exposure recovery assessment

To characterize recovery from ether anesthesia, four independent recovery trials were conducted using larvae from the same genetic crosses used in imaging experiments. In each trial, 10 second-instar larvae were exposed to diethyl ether vapor for 5 minutes at ∼25°C using the same Coplin chamber protocol described above (Section 2.4). After removal from the chamber, larvae were placed together on standard cornmeal-molassesagar media at 25°C and video-recorded for a minimum of 30 minutes. Two recovery endpoints were scored post hoc from recorded video: (1) twitch onset, defined as the first visible body wall movement, and (2) full movement, defined as the resumption of sustained forward locomotion. Larvae that did not reach a given endpoint within 30 minutes were recorded as right-censored at 30 minutes. Summary statistics (median, mean, range) were computed from uncensored observations only and therefore represent lower bounds on true population recovery times.

### 2.6 Imaging system

All images were acquired using a custom-built wide-field epifluorescence microscope (Fig. 3) constructed on a Thorlabs 30 mm cage system with a cage cube as the central junction for excitation and emission paths. The illumination path consisted of a 470 nm mounted LED (Thorlabs M470L4) powered by a Thorlabs LEDD1B LED driver (1200 mA maximum, with the current limit set to 1000 mA to match the LED maximum rating), with a GFP excitation bandpass filter (CWL 479 nm, BW 40 nm), collimated through an achromatic doublet lens (*f* = 100 mm) and relayed through a second 100 mm lens (L3) to the objective back focal plane. For GFP movement recordings, the LED was operated at the first major increment above zero; for calcium recordings, the LED was operated at maximum output. Excitation and emission were separated by a GFP dichroic mirror (cutoff 500 nm; Thorlabs MD498). Emission passed through a GFP bandpass filter (CWL 525 nm, BW 39 nm; Thorlabs MF525-39) and was focused by a 200 mm achromatic tube lens onto the camera sensor (Hamamatsu ORCA-Flash 4.0 V2, sCMOS, 6.5 µm pixel pitch, operated with 2 × 2 binning, yielding 1024 × 1024 pixels with an effective pixel size of 13.0 µm at the sensor). The objective was a Mitutoyo M Plan Apo 50× (NA 0.55, infinity-corrected, *f* = 200 mm), mounted via a Thorlabs SM1A12 adapter. With the 200 mm tube lens, the system magnification was 50×, giving a sample pixel size of 0.260 µm*/*pixel and a field of view of 266 × 266 µm. Images were acquired using HCImage software (Hamamatsu Photonics, DCAM drivers) at 16-bit depth. Acquisition parameters for both recording types are summarized in Table 1.

**Figure 3:**
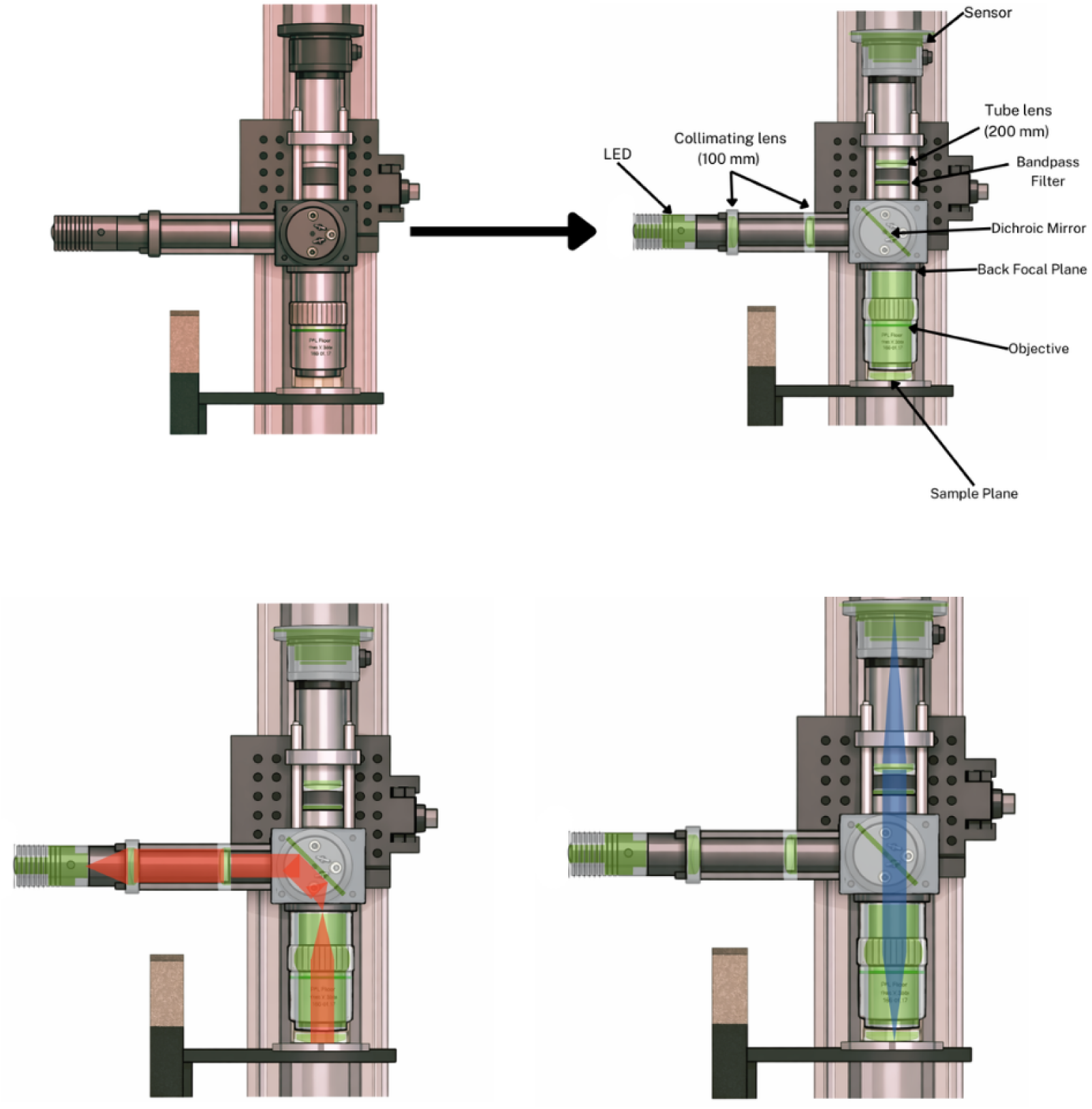
Wide-field epifluorescence microscope optical layout. Schematic showing illumination and detection beam paths, component positions, and optical layout. Component specifications detailed in Section 2.6.

**Table 1:** Acquisition parameters.

| Parameter | Movement (GFP) | Calcium (Neuro) |
| --- | --- | --- |
| Frame rate | 1 Hz | 33.33 Hz |
| Recording duration | 60 minutes | 5 minutes |
| Frames per recording | 3,600 | 10,000 |
| Exposure time | 1000 ms | 30 ms |
| Bit depth | 16-bit | 16-bit |
| File format | AVI (.avi) | TIFF (.tif) |

### 2.7 Motion correction and quantification

All motion analysis was performed using custom MATLAB software (R2024b, MathWorks). Frame-by-frame motion estimation used NoRMCorre [Pnevmatikakis and Giovannucci, 2017], a template-matching algorithm that computes rigid (translation-only) shifts by maximizing the correlation between each frame and a running template. Registration was performed with an initial search window of 30% of the field of view, with adaptive expansion up to 50% if motion exceeded the initial range. The algorithm processed data in 600-frame chunks with a memory target of 2 GB. Output per-frame *x*- and *y*-shift values were converted to micrometers using the calibrated pixel size (0.260 µm*/*pixel). Sign convention: positive *dx* indicates rightward sample displacement, positive *dy* indicates upward displacement (Cartesian convention). Motion correction applied corrective shifts via MATLAB’s `imtranslate(frame`, [−*dx*, +*dy*]), where the asymmetric signs reflect the conversion between the Cartesian motion coordinate system (*y*-up) and MATLAB’s image coordinate system (*y*-down).

#### Motion metrics

Three core metrics were computed from the shift time series for statistical comparison. Mean speed (µm*/*s) was defined as the total cumulative path length divided by recording duration, capturing the overall activity level. Median step size (µm) was the median of all frame-to-frame Euclidean displacements, providing a robust measure of typical per-frame motion. Jitter ratio (dimensionless) was defined as total path length divided by net displacement (the straight-line distance between first and last positions after median filtering with a 61-frame window, corresponding to approximately 1 minute at 1 Hz), characterizing movement pattern quality: values near 1 indicate directional drift while higher values indicate random or oscillatory motion.

### 2.8 Quality control system

Automated quality control flags were computed for each recording. For calcium data (`analyze_motion_and_QC_neuro.m`), MOVEMENT VALID required: registration failure rate (nan_frac) *<* 5%, fraction of frames at the search window boundary (`pegged_frac`) *<* 20%, no flatline detection (near-zero motion variance), and no unstable tracking. For GFP movement data (`analyze_motion_and_QC.m`), the equivalent tracking validity flag (`QC_PASS_tracking`) used stricter thresholds: `nan_frac` *<* 1% and `pegged_frac` *<* 1%. ROI ELIGIBLE, computed only for calcium recordings, additionally required that the total excursion not exceed 40% of the field of view in either axis (`boundary_exit` = false). Only MOVEMENT VALID recordings were included in motion comparisons; only ROI ELIGIBLE recordings entered the calcium imaging pipeline. This dual-flag system permits inclusion of recordings where motion tracking is reliable but the brain partially exited the field of view, which is informative for motion quantification but unsuitable for ROI extraction.

### 2.9 Calcium imaging processing pipeline

Calcium imaging analysis used a custom MATLAB pipeline (`COMPLETE_PIPELINE_RUN_ME.m`). Each ROI ELIGIBLE file was processed through the following steps: (1) loading precomputed shifts from the motion QC output; (2) applying rigid motion correction by translating each frame by the negative of the estimated shift; (3) saving the corrected stack in BigTIFF format; (4) generating a brain mask using thresholding and morphological operations; (5) computing local correlation (*C_n_*) and peak-to-noise ratio (PNR) images; (6) detecting candidate ROI seeds as local maxima in the *C_n_* ×PNR product that exceed an adaptive threshold; (7) growing ROI regions from seeds using correlation-based expansion; (8) extracting raw fluorescence traces and estimating neuropil contamination from annular regions surrounding each ROI; (9) computing neuropil-corrected Δ*F/F* using a sliding baseline; (10) detecting calcium events using physiologically constrained peak finding; and (11) computing per-ROI quality metrics.

ROI detection used adaptive thresholding based on *C_n_* and PNR maps, following approaches described for CaImAn [Giovannucci et al., 2019]. The expected neuron radius (`gSig`) was set to 3 pixels (0.78 µm), with a minimum local correlation threshold of 0.3 and a minimum PNR of 2.0. ROI area was constrained to 12–120 pixels^2^ (0.81–8.1 µm^2^), corresponding to equivalent circular diameters of approximately 1.0–3.2 µm, appropriate for second-instar *Drosophila* neuron somata. Candidate ROIs with more than 30% of their area overlapping previously placed ROIs were excluded rather than merged. After initial detection, signal quality was assessed for each recording (`Assess_ROI_Quality.m`) to evaluate pairwise synchrony across ROIs as an indicator of shared neuropil contamination. Samples were categorized based on mean pairwise synchrony: clean signal (*<* 0.15), mild neuropil contamination (0.15–0.30), high neuropil (0.30–0.50), severe neuropil (*>* 0.50), or artifact (extreme Δ*F/F* values exceeding 5.0 or below −2.0). These categories are reported for transparency but were not used as an exclusion gate; all ROI ELIGIBLE recordings proceeded to refinement regardless of quality category. Recordings passing quality assessment then entered a multi-criteria refinement step (`ROI_REFINEMENT.m`) that sequentially evaluated each ROI against ten criteria: (1) extreme SNR values exceeding the physiological range for wide-field imaging (SNR *>* 150); (2) edge contact, where ROIs with more than 20% of their perimeter touching the brain mask boundary were removed; (3) morphological shape, enforcing minimum compactness (4*πA/P* ^2^ ≥ 0.20), bounding box extent (0.35–0.95), aspect ratio (≤ 3.0), solidity (≥ 0.55), and eccentricity (≤ 0.93); (4) negative transients, removing ROIs with dips exceeding 3 SD below baseline; (5) step artifacts, removing ROIs with single-frame Δ*F/F* jumps exceeding 0.5; (6) non-physiological rise/decay kinetics inconsistent with GCaMP8f dynamics (rise/decay ratio *>* 3.0, decay time constant outside 0.05–8.0 s); (7) sustained plateaus, where signal remained above 70% of peak amplitude for more than 20 s without decay; (8) pairwise trace correlation exceeding 0.7, with the lower-SNR member of each correlated pair removed; (9) spatial proximity, where the lower-SNR member of ROI pairs with centers closer than 5 pixels was removed; and (10) insufficient activity (zero detected events) or baseline drift. The ROI counts, SNR values, and event rates reported in the Results (Table 5) reflect the refined dataset. A breakdown of removal counts by criterion is shown in Figure 10. After detection, each ROI was assigned a signal-to-noise ratio (SNR), defined as peak Δ*F/F* amplitude divided by the standard deviation of the baseline fluorescence. ROIs with SNR *<* 2.0 were excluded from downstream analysis. An adaptive relaxation for recordings with uniformly low signal was implemented but was not triggered for any recording in this dataset.

Neuropil correction used an annular region constructed by morphological dilation of each ROI mask. The inner boundary was defined by dilating the ROI mask with a disk structuring element of radius equal to `gSig` (3 pixels), and the outer boundary by dilating with a disk of radius 2.5 × `gSig` (8 pixels), with a minimum annulus width of 4 pixels enforced. Pixels belonging to other ROIs were excluded from each neuropil annulus. Corrected fluorescence was computed as *F*_corr_(*t*) = *F*_raw_(*t*) − 0.7 × *F*_neuropil_(*t*), where 0.7 is a widely used neuropil correction coefficient adopted in both two-photon and wide-field calcium imaging studies [Chen et al., 2013, Dana et al., 2019]. Baseline fluorescence was estimated using a sliding minimum filter (30-second window) followed by moving average smoothing of equal window length [Jia et al., 2011]. Prior to event detection, neuropil-corrected Δ*F/F* traces were smoothed using a Savitzky–Golay filter (3rd-order polynomial, 5-frame window corresponding to approximately 150 ms at 33.33 Hz) to reduce high-frequency noise while preserving transient shape.

Event detection used MATLAB’s `findpeaks`() with constraints optimized for GCaMP8f kinetics: minimum prominence of 0.03 Δ*F/F* (3%), minimum peak width of 200 ms (∼7 frames at 33.33 Hz), maximum peak width of 10 s, minimum peak distance of 300 ms, and minimum rise rate of 0.01 Δ*F/F* per second. Events failing the rise rate criterion were excluded to reject slow drift artifacts.

#### Global signal regression

To address shared neuropil contamination that affects the entire field of view [Musall et al., 2019], the pipeline includes an optional global signal regression step. The global signal was computed from the mean fluorescence of all brain mask pixels excluding ROI territories, sampled at 500 evenly spaced frames and interpolated to full temporal resolution using piecewise cubic Hermite interpolation. This signal was converted to Δ*F/F* scale and regressed from each ROI trace using ordinary least-squares regression with an intercept. This approach estimates and removes the component of each ROI’s trace that is linearly predicted by the global neuropil signal, while preserving ROI-specific dynamics. Pairwise synchrony (mean absolute pairwise Pearson correlation between all ROI trace pairs) was computed before and after regression to verify that shared variance was reduced without over-correction. All calcium imaging results reported in this paper were computed from traces after global signal regression.

### 2.10 Statistical analysis

Group comparisons for motion metrics used three complementary tests per metric: (1) a two-sided permutation test (*B* = 10,000) as the primary hypothesis test, appropriate for small samples with unknown distributions; (2) Welch’s *t*-test as a parametric reference; and (3) the Mann–Whitney *U* test as a non-parametric rank-based comparison. *P*-values from permutation tests were corrected for multiple comparisons using the Benjamini– Hochberg false discovery rate (FDR) procedure [Benjamini and Hochberg, 1995]. Adjusted *p*-values (*q*-values) are reported with a significance threshold of *q <* 0.05.

Effect sizes were quantified using Hedges’ *g*, a bias-corrected standardized mean difference [Hedges, 1981]. Effects were interpreted following conventional thresholds as negligible (|*g*| *<* 0.2), small (0.2–0.5), medium (0.5–0.8), or large (≥ 0.8) [Cohen, 1988]. Ninety-five percent confidence intervals for mean differences and for Hedges’ *g* were computed via bootstrap resampling (*B* = 10,000) [Efron and Tibshirani, 1993]. All analyses used fixed random seeds for reproducibility: `rng(42, ‘twister’)` for the calcium imaging pipeline and statistical comparison scripts, and `rng(0, ‘twister’)` for the motion tracking and QC scripts.

Two analysis windows were examined for movement data: full recording duration (∼60 minutes) as the primary analysis, and the first 30 minutes as a secondary analysis. The 30-minute window was selected a priori based on the hypothesis that anesthetic effects may decay over time, but is treated as exploratory because it was not the primary endpoint.

Calcium imaging group comparisons used Cliff’s delta (*δ*; Cliff 1993) as a non-parametric effect size measure with bootstrap 95% confidence intervals, applied at the ROI level for SNR and event rate. Multiple comparisons across calcium imaging metrics were corrected using the Bonferroni procedure.

#### Sample flow

For movement experiments, 35 larvae were prepared across both conditions (19 control, 16 experimental). Four control larvae were excluded: two for first-instar staging, one for third-instar staging, and one for ambient light contamination during imaging. One experimental larva was excluded for first-instar staging. This yielded 15 analyzable recordings per condition. All 30 included recordings passed automated QC (MOVEMENT VALID). For calcium experiments, 52 larvae were prepared (24 control, 28 experimental). One control larva was excluded for first-instar staging. Six experimental larvae were excluded for incorrect instar staging (3 first-instar, 3 third-instar), and one additional experimental recording (S11) failed automated QC due to excessive motion, yielding 23 control and 21 experimental recordings classified as ROI ELIGIBLE.

#### Randomization and blinding

Animals were assigned to experimental conditions opportunistically within each imaging session based on available larvae. No systematic bias was applied during selection, and downstream analysis was performed using automated pipelines applied identically across conditions. Statistical comparisons were performed on files identified only by condition label, without access to individual animal identifiers during analysis.

## 3 Results

### 3.1 Multimodal immobilization reduces motion across all metrics

We compared immobilization effectiveness between hydrogel-only (Control, *N* = 15) and diethyl ether + hydrogel (Experimental, *N* = 15) preparations using three non-redundant motion metrics derived from frame-by-frame shift estimates over ∼60-minute wide-field GFP recordings. All recordings in both groups passed automated QC (MOVEMENT VALID = true), with no tracking failures, search ceiling saturation, or flatline events detected.

Across the full recording duration, the multimodal condition significantly reduced motion on all three core metrics after FDR correction (Table 2; Fig. 4). Mean speed was reduced by 90.7% (mean difference −0.395 µm*/*s, 95% CI [−0.653, −0.217]; Hedges’ *g* = −1.18; *q <* 0.001). Median step size was reduced by 84.5% (mean difference −0.141 µm, 95% CI [−0.219, −0.081]; *g* = −1.36; *q <* 0.001). Jitter ratio was reduced by 88.7% (mean difference −88.14, 95% CI [−175.0, −29.26]; *g* = −0.80; *q <* 0.001). All confidence intervals for the mean difference excluded zero.

**Figure 4:**
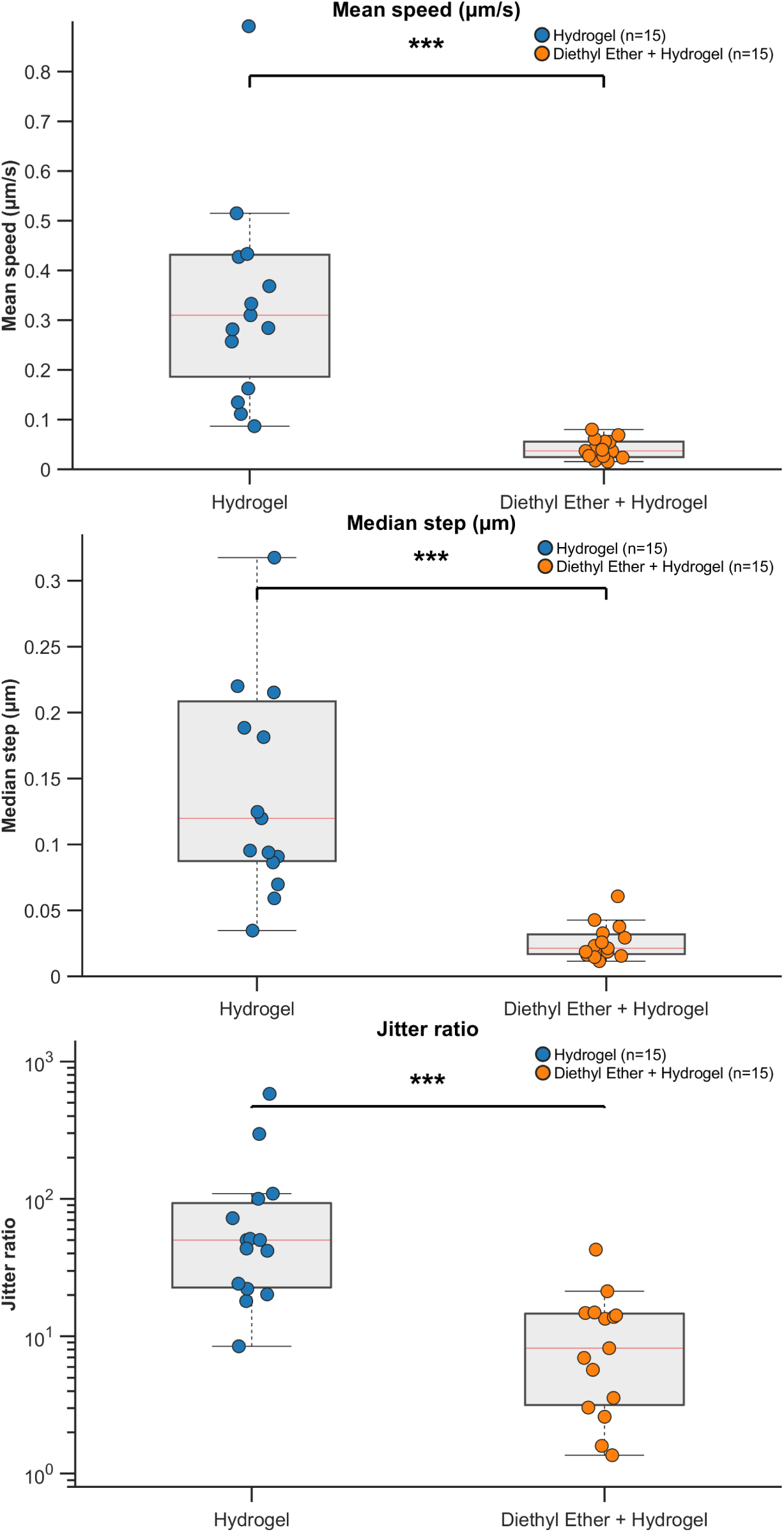
Full-duration motion comparison between immobilization conditions. Box plots with individual data points for (A) mean speed (µm*/*s), (B) median step (µm), and (C) jitter ratio over the full ∼60-minute recording. Blue = Control (hydrogel only, *N* = 15); Orange = Experimental (ether + hydrogel, *N* = 15). Boxes show median and interquartile range; whiskers extend to 1.5× IQR. \*\*\**q <* 0.001.

**Table 2:** Full-duration motion comparison (Experimental − Control). Negative effects indicate reduced motion in the Experimental group. 95% CIs are for mean differences (bootstrap, *B* = 10,000). *p*_perm_ = permutation test (*B* = 10,000); *q*_FDR_ = Benjamini– Hochberg adjusted *p*-value.

| Metric | Mean diff | 95% CI | Hedges' $g$ | % change | $p_{\text{perm}}$ | $q_{\text{FDR}}$ |
| --- | --- | --- | --- | --- | --- | --- |
| Mean speed ($\mu\text{m/s}$) | $-0.395$ | $[-0.653, -0.217]$ | $-1.18$ | $-90.7\%$ | $< 0.001$ | $< 0.001$ |
| Median step ($\mu\text{m}$) | $-0.141$ | $[-0.219, -0.081]$ | $-1.36$ | $-84.5\%$ | $< 0.001$ | $< 0.001$ |
| Jitter ratio | $-88.14$ | $[-175.0, -29.26]$ | $-0.80$ | $-88.7\%$ | $< 0.001$ | $< 0.001$ |

### 3.2 Stronger effects in the first 30 minutes of recording

As a secondary analysis, we examined the first 30 minutes of each recording separately. This earlier window targets the period of expected maximal chemical suppression (see Section 2.10). Effects were consistently stronger in this earlier window (Table 3; Fig. 5). All three core metrics reached FDR-corrected significance with uniformly large effect sizes: mean speed (*g* = −1.23, 93.7% reduction, *q <* 0.001), median step (*g* = −1.51, 92.6% reduction, *q <* 0.001), and jitter ratio (*g* = −1.10, 81.3% reduction, *q <* 0.001).

**Figure 5:**
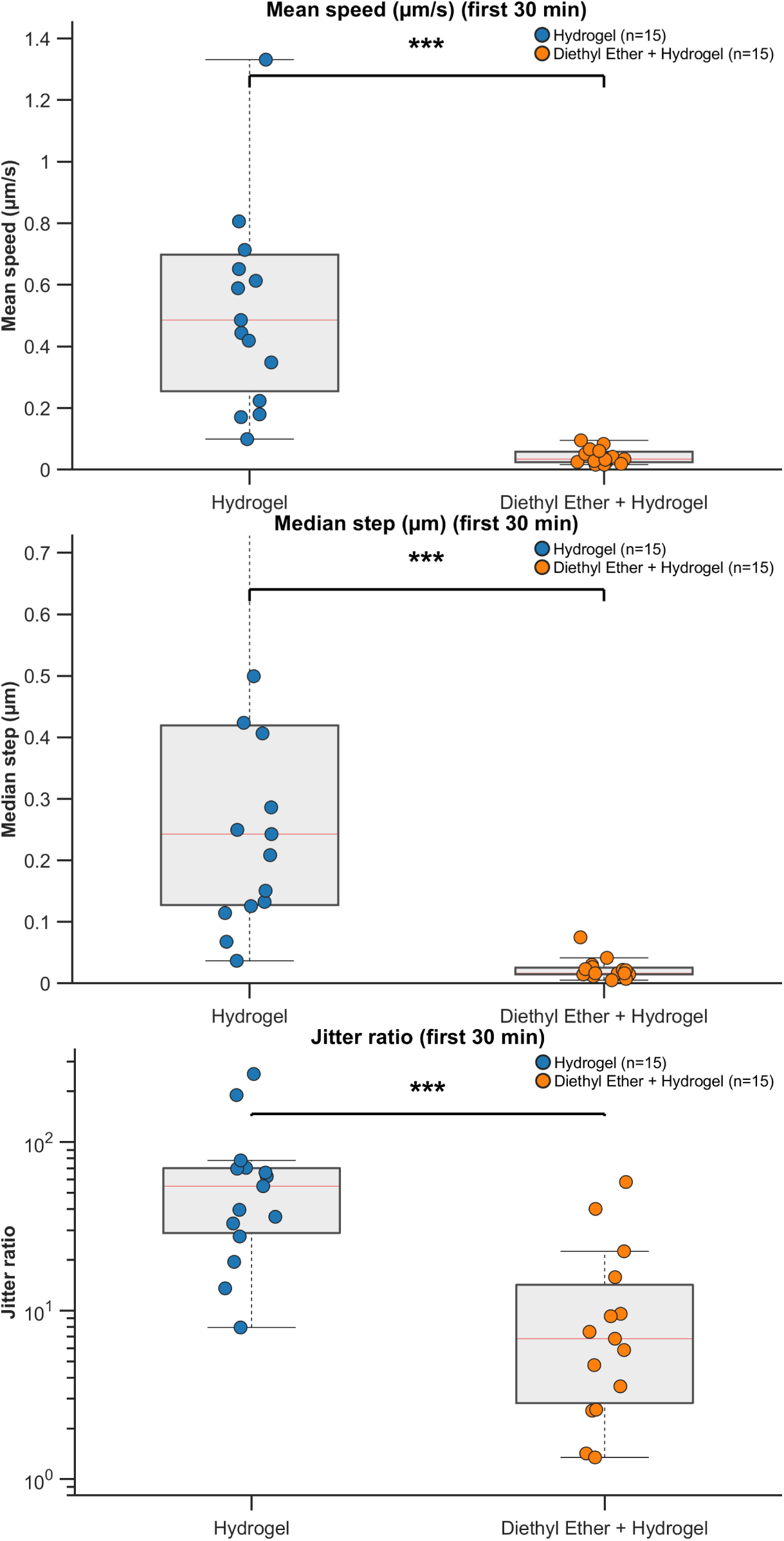
Motion comparison in the first 30 minutes of recording (exploratory analysis). Box plots with individual data points for (A) mean speed, (B) median step, and (C) jitter ratio. Conventions as in Figure 4. \*\*\**q <* 0.001.

**Table 3:** First-30-minute motion comparison (exploratory). Conventions as in. **Table 2**.

| Metric | Mean diff | 95% CI | Hedges' $g$ | % change | $p_{\text{perm}}$ | $q_{\text{FDR}}$ |
| --- | --- | --- | --- | --- | --- | --- |
| Mean speed ($\mu\text{m/s}$) | -0.624 | [-1.01, -0.350] | -1.23 | -93.7% | < 0.001 | < 0.001 |
| Median step ($\mu\text{m}$) | -0.288 | [-0.426, -0.171] | -1.51 | -92.6% | < 0.001 | < 0.001 |
| Jitter ratio | -55.39 | [-91.60, -25.70] | -1.10 | -81.3% | < 0.001 | < 0.001 |

### 3.3 Temporal dynamics of immobilization

Comparing effect sizes between analysis windows reveals the temporal dynamics of immobilization (Table 4). Several metrics showed notably larger effects in the first 30 minutes: jitter ratio increased from *g* = −0.80 to *g* = −1.10, with the largest relative change (Δ*g* = −0.30) among the three metrics. The time course overlay of per-minute median step size (Fig. 6) shows that the experimental group maintains consistently lower motion throughout the recording, with the largest separation during the initial portion and a gradual decline in control group motion over time before it reaches a lower plateau. This pattern is consistent with the hypothesis that ether provides stronger initial chemical suppression that partially wanes, while the hydrogel maintains mechanical restraint throughout.

**Figure 6:**
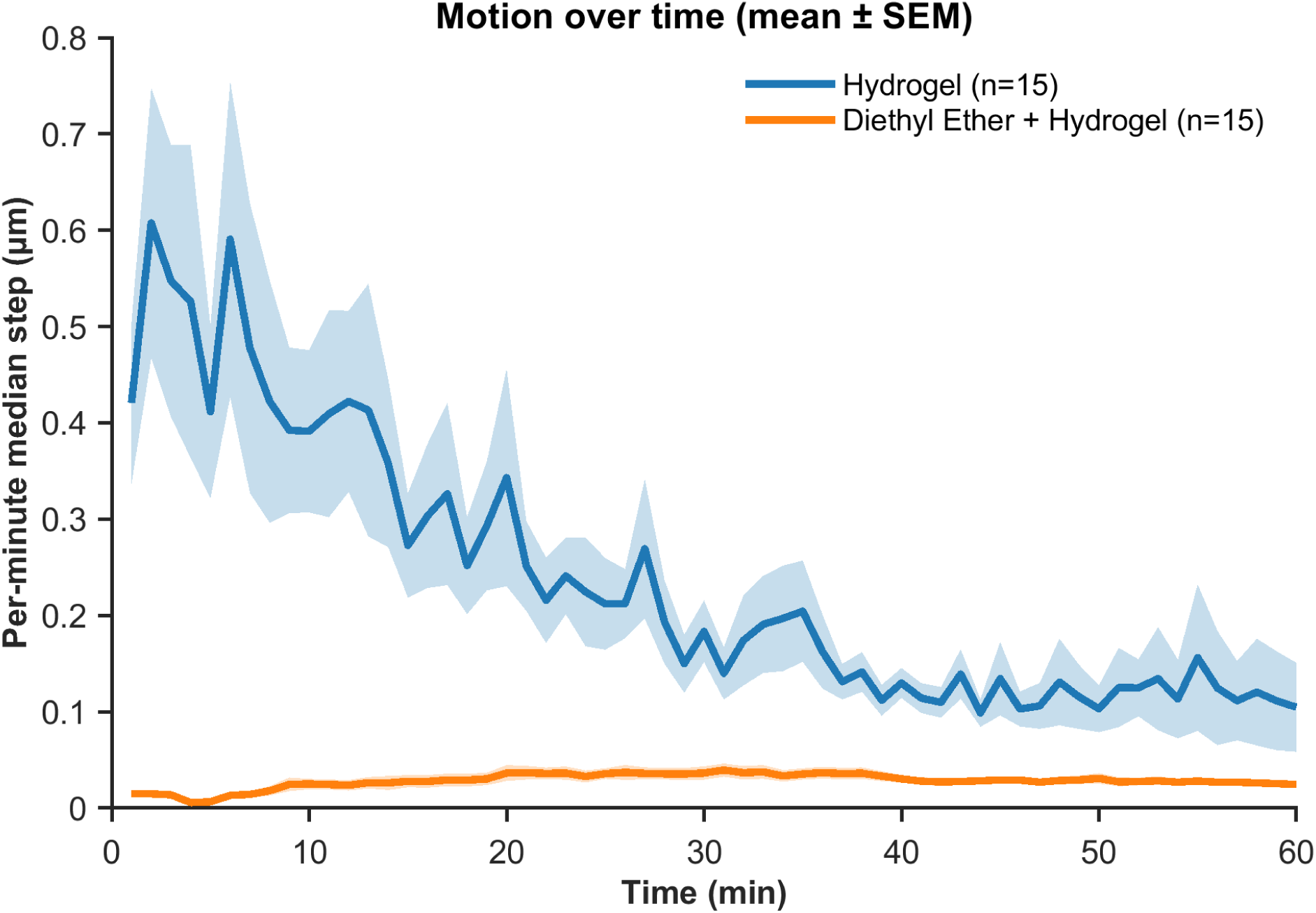
Temporal dynamics of motion by condition. Mean ± SEM per-minute median step (µm) plotted over the recording duration for Control (blue) and Experimental (orange) groups. Shaded regions indicate SEM.

**Table 4:**
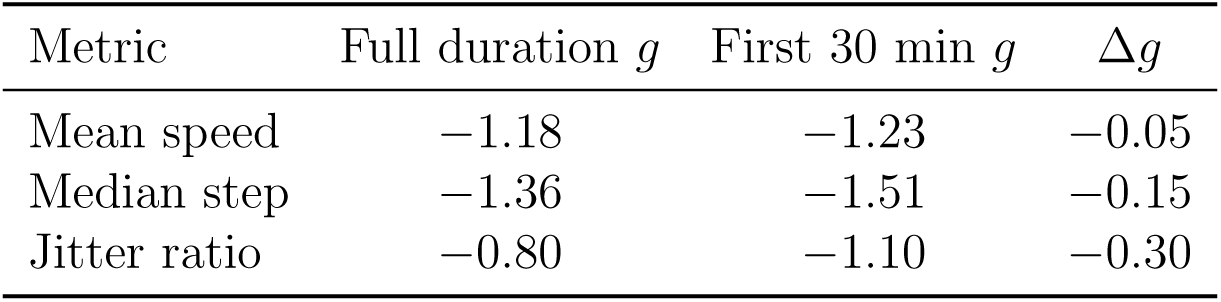
Comparison of effect sizes (Hedges’ *g*) between full-duration and first-30-minute analysis windows. Δ*g* = first 30 min *g* − full duration *g*; more negative values indicate stronger early effects.

### 3.4 Calcium imaging: quality control and eligibility gating

For calcium recordings (33.33 Hz, ∼5 minutes), the dual-flag QC system identified recordings suitable for ROI extraction. In the control group, 23 of 24 total recordings were classified as `ROI_ELIGIBLE` (1 excluded for first-instar staging). In the experimental group, 22 recordings entered the QC pipeline after staging exclusions; of these, one recording (S11) failed both `MOVEMENT_VALID` and `ROI_ELIGIBLE` flags due to excessive motion, yielding 21 `ROI_ELIGIBLE` experimental recordings.

Post-pipeline signal quality assessment revealed an asymmetry in neuropil contamination between conditions. In the control group, 8 of 23 recordings (34.8%) were categorized as having acceptable signal quality (clean or mild neuropil contamination) based on pairwise synchrony thresholds, compared to 16 of 21 (76.2%) in the experimental group. The higher exclusion rate in controls likely reflects greater residual motion, which increases shared fluorescence fluctuations across ROIs and elevates pairwise synchrony scores above the quality thresholds. Because ROI-level refinement addresses these contamination sources on a per-ROI basis, all `ROI_ELIGIBLE` recordings were retained for downstream analysis rather than applying sample-level exclusions.

### 3.5 Calcium imaging results

Both conditions yielded successful ROI detection and identification of calcium transients (Table 5; Figs. 9, 7, 8). Control recordings (*N* = 23) produced 368 total ROIs (mean 16.0 ± 15.9 per sample) with high signal-to-noise ratio (SNR 30.4 ± 16.9) and a mean event rate of 0.228 ± 0.113 Hz. Experimental recordings (*N* = 21) produced 295 total ROIs (mean 14.0 ± 17.4 per sample) with lower but still substantial SNR (18.0 ± 10.6) and a slightly higher mean event rate of 0.309 ± 0.188 Hz.

**Table 5:** Calcium imaging summary by condition.

| Metric | Control ($N = 23$) | Experimental ($N = 21$) |
| --- | --- | --- |
| Total ROIs | 368 | 295 |
| ROIs per sample (mean ± SD) | $16.0 \pm 15.9$ | $14.0 \pm 17.4$ |
| Mean SNR (mean ± SD) | $30.4 \pm 16.9$ | $18.0 \pm 10.6$ |
| Event rate, Hz (mean ± SD) | $0.228 \pm 0.113$ | $0.309 \pm 0.188$ |

**Figure 7:**
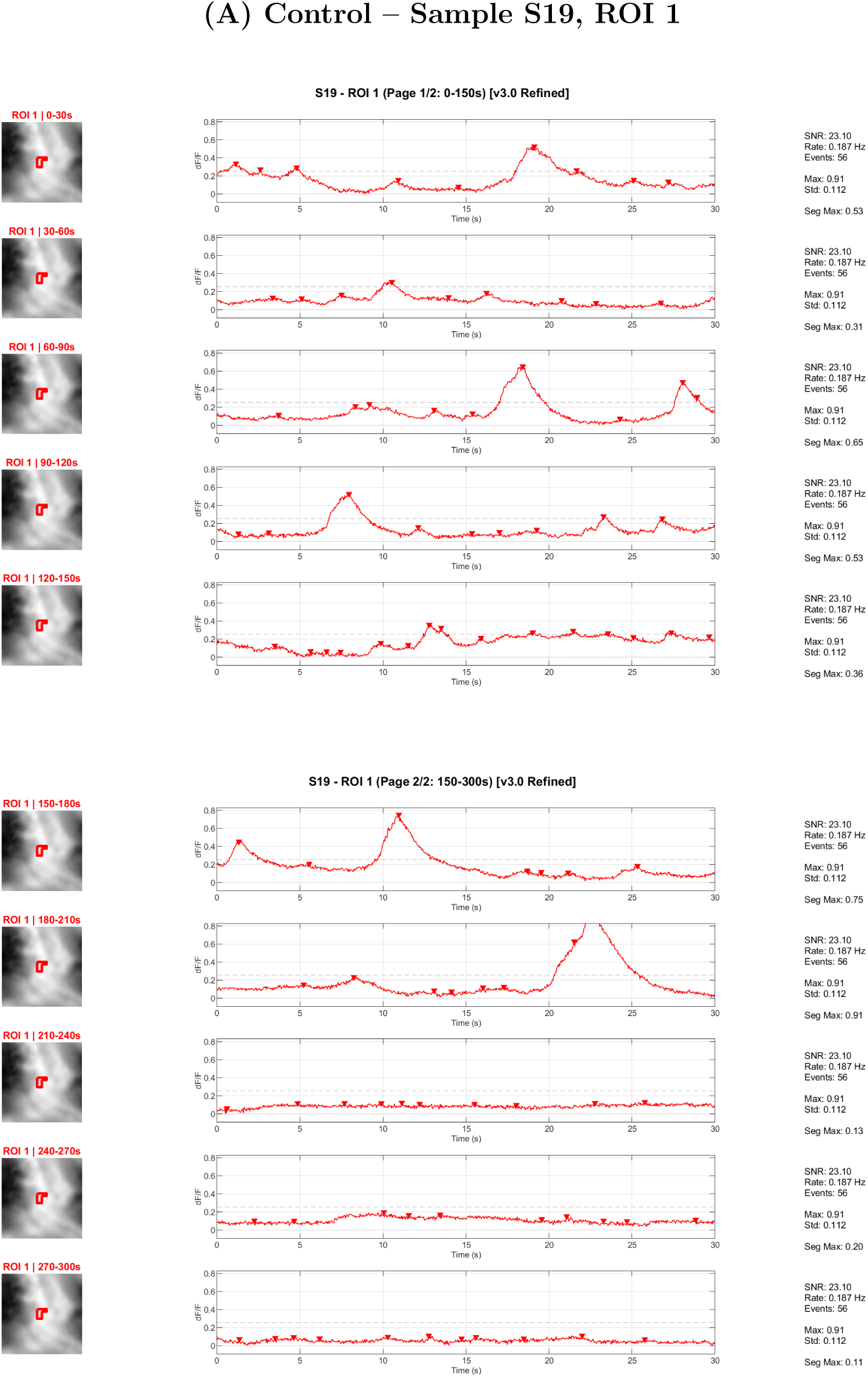
Representative calcium traces from a control recording (sample S19, ROI 1; SNR = 23.1, event rate = 0.187 Hz). Neuropil-corrected Δ*F/F* traces shown in 30-second segments across the full ∼5-minute recording. Red triangles indicate detected calcium events. Dashed lines show the event detection threshold. The ROI spatial footprint is shown at left for each segment.

**Figure 8:**
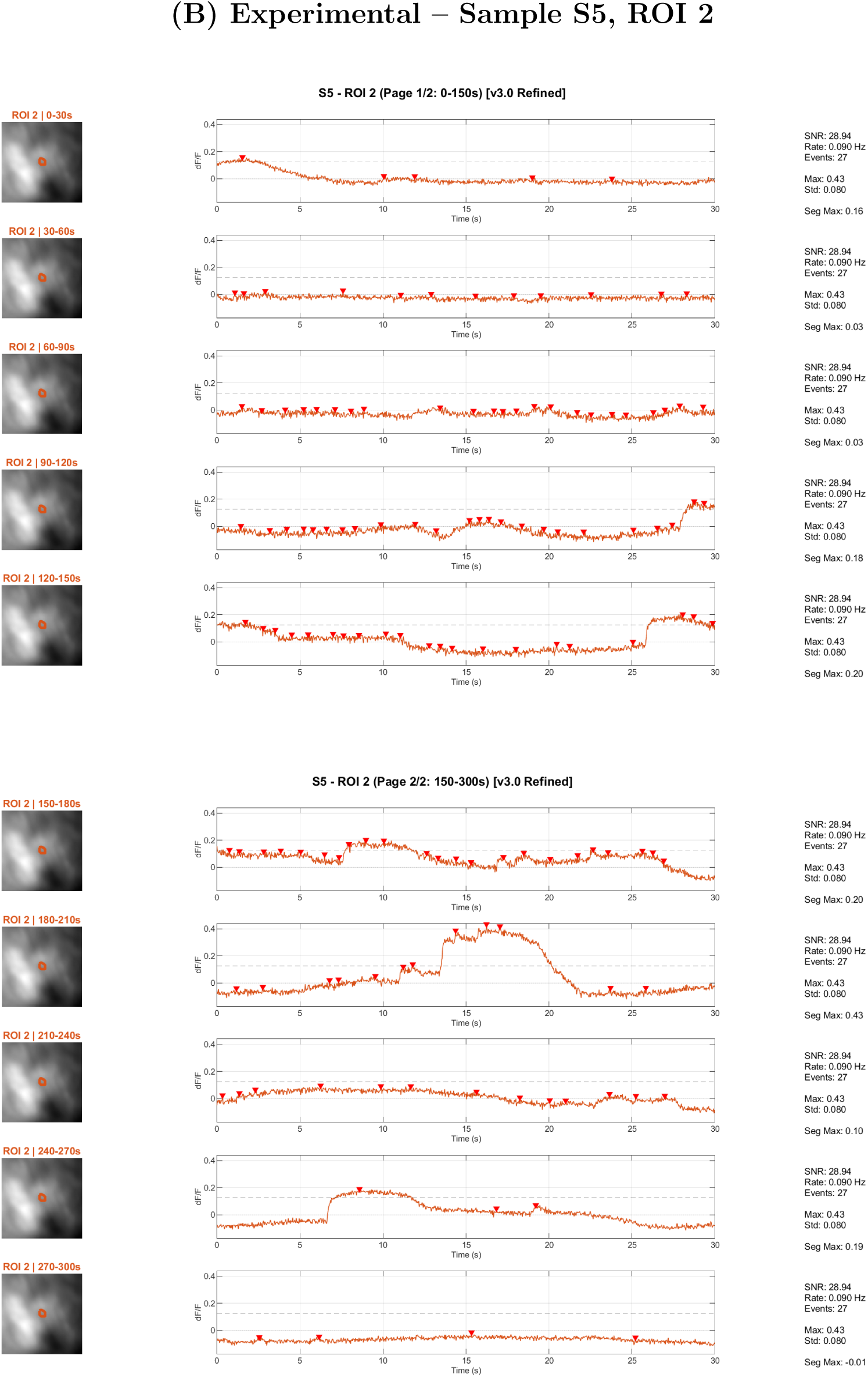
Representative calcium traces from an experimental recording (sample S5, ROI 2; SNR = 28.9, event rate = 0.090 Hz). Conventions as in Figure 7. This ROI illustrates that ether-treated preparations can produce high-SNR traces with clearly resolved calcium transients, consistent with detectable calcium dynamics under multimodal immobilization.

**Figure 9:**
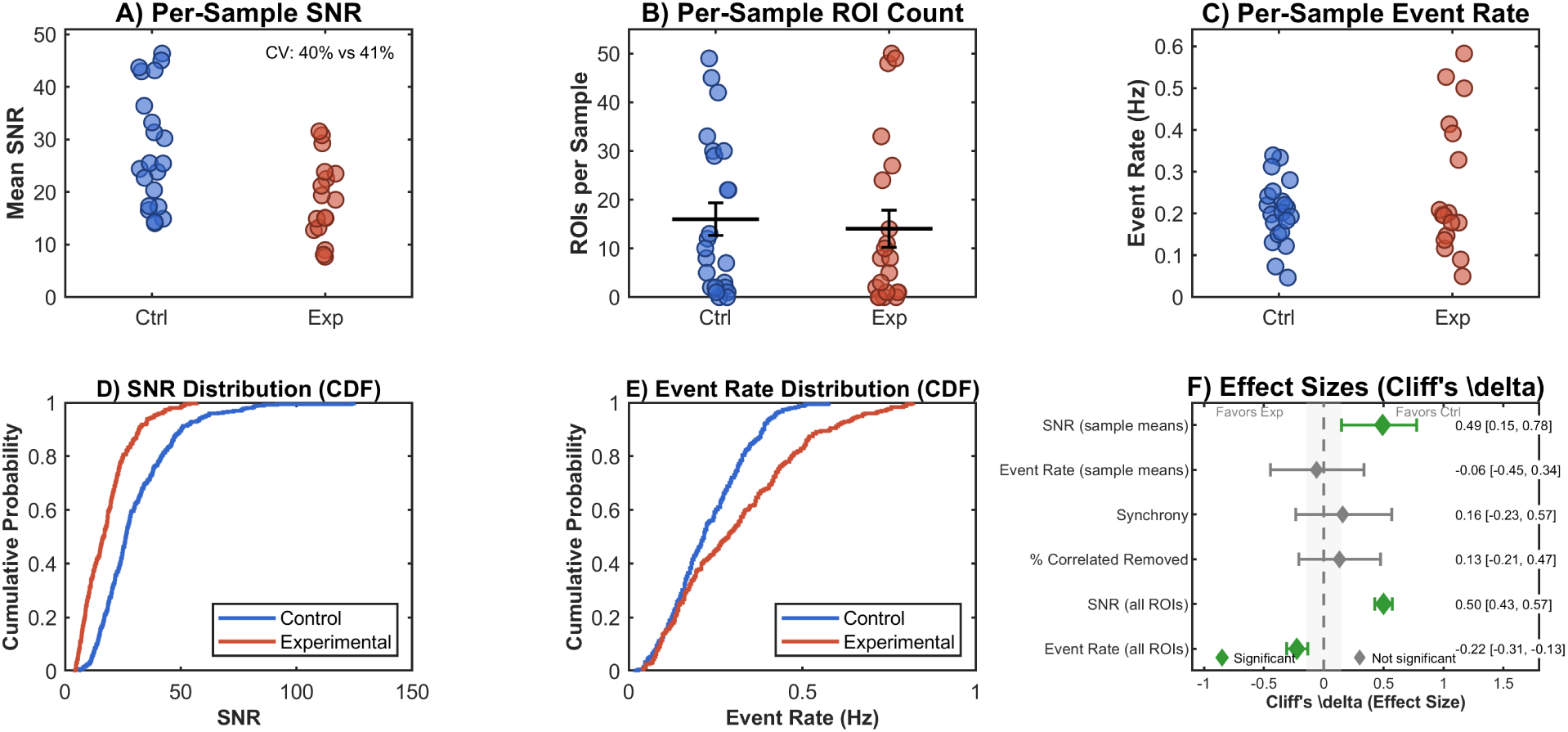
Calcium imaging group comparison. (A) Per-sample mean SNR, (B) Per-sample ROI count, (C) Per-sample event rate. (D) CDF of SNR across all ROIs. (E) CDF of event rate. (F) Forest plot of Cliff’s *δ* effect sizes with 95% bootstrap CIs.

**Figure 10:**
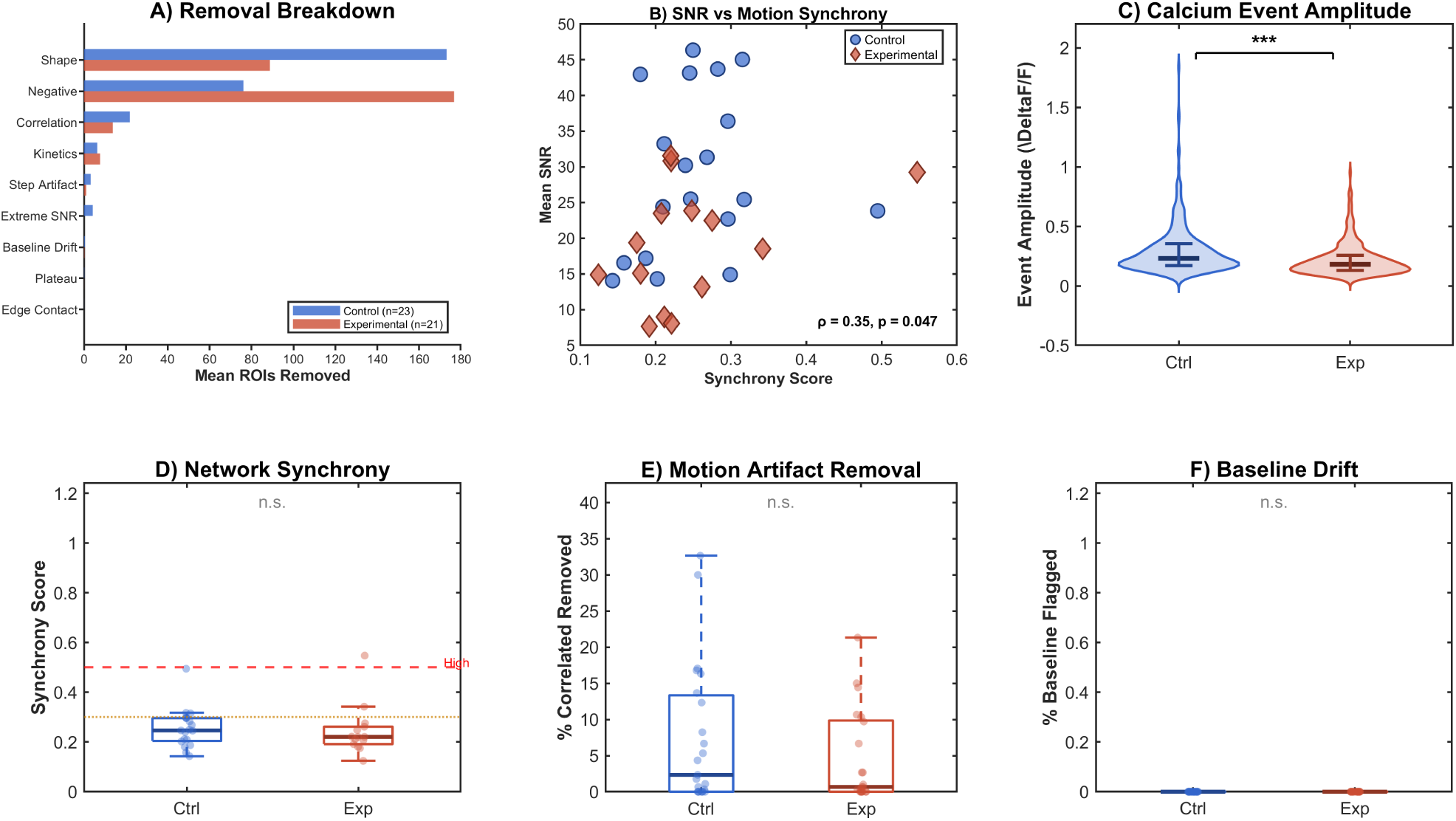
ROI quality assessment. (A) ROI removal breakdown by criterion. (B) SNR vs. motion synchrony score. (C) Calcium event amplitude by group. (D–F) Network synchrony, motion artifact removal rate, and baseline drift flag rate by group.

At the ROI level, SNR was significantly higher in control recordings (Cliff’s *δ* = 0.50, 95% CI [0.43, 0.57], *p <* 0.001), while event rate was modestly higher in the experimental group (*δ* = −0.22, 95% CI [−0.31, −0.13], *p <* 0.001). At the sample level, mean SNR remained higher in controls (*δ* = 0.49, *p* = 0.011) but did not survive Bonferroni correction (*p*_adj_ = 0.106), and sample-level event rate showed no significant difference (*δ* = −0.06, *p* = 0.77). Sample-level SNR and event rate comparisons include only samples with at least one retained ROI after refinement (*N* = 21 control, *N* = 17 experimental); the remaining 2 control and 4 experimental samples had all ROIs removed during the multi-criteria refinement process and therefore had no summary metrics to compare. ROI count per sample did not differ significantly between groups. Network synchrony and the proportion of correlated ROIs removed during refinement also showed no significant group differences, suggesting that the overall structure of detected neural activity is similar between conditions.

The lower SNR in experimental recordings could reflect several factors: ether-induced changes in baseline fluorescence, differences in tissue optical properties, or the relationship between motion reduction and signal quality. Importantly, the modestly higher event rate observed at the ROI level in the experimental group argues against complete neural suppression by ether. Rather, detected transients in ether-treated preparations are somewhat more frequent but of lower amplitude, consistent with altered but not abolished calcium dynamics. Event rates observed across both groups (0.14–0.77 Hz) fall within the biologically plausible range for immobilized larval preparations where locomotor CPG activity is suppressed.

### 3.6 Post-exposure recovery

Across four independent trials (*N* = 40 larvae total), 70.0% of larvae (28/40) exhibited twitching within 30 minutes of ether removal, and 60.0% (24/40) recovered full movement within the same window (Table 6). Among larvae that recovered, median twitch onset was 12.5 minutes (mean 12.8 ± 5.6 min; range 0.03–22.6 min) and median time to full movement was 17.7 minutes (mean 17.9 ± 6.1 min; range 0.12–25.9 min). One larva exhibited anomalously rapid recovery (twitch at 0.03 min, full movement at 0.12 min), likely reflecting incomplete initial anesthesia; this observation was retained but did not meaningfully influence summary statistics. Larvae that did not recover within the 30-minute observation window were not monitored further, so recovery beyond this time point was not assessed.

**Table 6:** Post-exposure recovery summary. Recovery assessed over a 30-minute observation window following 5 minutes of diethyl ether exposure at ∼25°C. Statistics computed from uncensored observations only.

| Metric | Twitch Onset | Full Movement |
| --- | --- | --- |
| Total observed | 40 | 40 |
| Recovered within 30 min | 28 (70.0%) | 24 (60.0%) |
| Median recovery time (min) | 12.5 | 17.7 |
| Mean ± SD (min) | $12.8 \pm 5.6$ | $17.9 \pm 6.1$ |
| Range (min) | 0.03–22.6 | 0.12–25.9 |

## 4 Discussion

### 4.1 Summary of findings

We evaluated a multimodal immobilization strategy combining PF-127 hydrogel with brief diethyl ether exposure for calcium imaging in second-instar *Drosophila* larvae. The combined approach produced significant reductions in residual motion compared to hydrogel alone, with large effect sizes observed for mean speed (*g* = −1.18) and median step size (*g* = −1.36) over the full recording duration. The effect was strongest during the first 30 minutes, during which all three core metrics reached FDR-corrected significance with uniformly large effect sizes (|*g*| = 1.10–1.51). Calcium imaging in both conditions yielded successful ROI detection with biologically plausible event rates, though experimental preparations showed lower SNR and modestly higher event rates at the ROI level compared to the control.

### 4.2 Mechanism of ether immobilization and temporal dynamics

Diethyl ether is a volatile general anesthetic whose mechanisms of action are multifaceted. In mammals, ether is thought to potentiate inhibitory GABA_A_ receptors and glycine receptors, inhibit excitatory NMDA receptors, and modulate two-pore-domain potassium channels, collectively depressing neural excitability and suppressing motor output [Hemmings et al., 2005]. In *Drosophila*, genetic evidence points to voltage-gated sodium channels as particularly important: mutations in the *para* sodium channel gene strongly modulate ether sensitivity, and this effect depends primarily on channel genotype rather than membrane properties, suggesting a direct molecular interaction between ether and the sodium channel [Tanaka and Gamo, 2001]. The *para* channel is the sole voltage-gated sodium channel in *Drosophila*, meaning that ether-induced sodium channel modulation could broadly affect neural excitability throughout the nervous system. While these mechanisms were not directly assessed in the present study, they provide context for interpreting the observed effects on motion and calcium dynamics.

The temporal decay of immobilization we observed, with stronger effects in the first 30 minutes and persistent but reduced effects over 60 minutes, is consistent with ether’s pharmacokinetics. As a highly volatile compound, absorbed ether is expected to dissipate rapidly from larval tissue once the animal is removed from the vapor chamber, and its anesthetic effects would diminish as the tissue concentration decreases. The fact that motion remains substantially lower in the experimental group even at 60 minutes likely reflects the continued mechanical restraint provided by the hydrogel matrix, which maintains physical immobilization after chemical suppression has partially worn off. This complementary mechanism is the central rationale for the multimodal approach.

### 4.3 Implications of calcium imaging results

The calcium imaging results reveal a nuanced picture. The higher SNR in control recordings (Cliff’s *δ* = 0.50) was unexpected, since one might predict that reduced motion in the experimental group would improve signal quality. Several factors may explain this. Ether could alter baseline fluorescence through effects on intracellular calcium homeostasis or pH-dependent changes in GCaMP fluorescence. Additionally, if ether partially suppresses neural activity, the peak calcium transients that define the “signal” in the SNR calculation would be smaller, lowering the ratio even if baseline noise is also reduced.

The modestly higher event rate in experimental preparations at the ROI level (*δ* = −0.22) is notable because it argues against the concern that ether exposure simply abolishes neural activity, though this difference did not reach significance at the sample level after correction for multiple comparisons. Several interpretations are possible. Reduced motion in the experimental group may improve the reliability of event detection by reducing motion-correlated fluorescence fluctuations that could mask real transients. Alternatively, ether may alter the balance between excitation and inhibition in ways that produce more frequent but lower-amplitude calcium events. The fact that network synchrony did not differ significantly between groups further suggests that the overall pattern of coordinated neural activity is not substantially disrupted.

These findings have practical implications for experimental design. For studies focused on population-level activity patterns, connectivity, or event timing, the multimodal approach may retain sufficient features of neural dynamics while substantially improving preparation stability. For studies requiring maximal signal amplitude or minimal pharmacological perturbation, hydrogel-only immobilization may be preferable despite higher residual motion. We recommend that users explicitly report which immobilization method was used and present motion metrics alongside calcium imaging results to enable informed interpretation.

### 4.4 Comparison to existing methods

Compared to microfluidic devices, our method requires no specialized fabrication, clean-room access, or engineering expertise. The entire protocol can be learned in a single session using standard laboratory supplies. While microfluidic platforms may provide more complete immobilization, they impose substantially higher technical and logistical barriers that limit their adoption, particularly in laboratories focused on genetics or neuroscience rather than microengineering. The multimodal approach described here offers a practical middle ground: significantly better immobilization than hydrogel alone, with no increase in required infrastructure.

Compared to protocols relying on prolonged anesthesia, our approach uses a single 5-minute ether exposure followed by mechanical maintenance of immobilization, whereas Kakanj et al. [2020] demonstrated that 3–4.5 minutes of ether alone can sustain immobilization for up to 8 hours without additional restraint. The addition of hydrogel mechanical support in our protocol is intended to extend effective immobilization as chemical effects partially wane, while keeping the total anesthetic exposure brief. Compared to cold anesthesia, our protocol confines any temperature perturbation to the pre-imaging phase, with all data acquisition occurring at room temperature.

Recovery trials confirmed that larvae resume movement after the same ether exposure protocol used for imaging. Median twitch onset occurred at 12.5 minutes and full movement at 17.7 minutes, with 70% and 60% of larvae reaching these endpoints within 30 minutes, respectively. The 40% of larvae that did not recover full movement within the observation window is consistent with the prolonged immobilization reported by Kakanj et al. [2020], who observed that 3–4.5 minutes of ether exposure can sustain immobilization for up to 8 hours. In our protocol, continued immobilization beyond 30 minutes is advantageous for imaging purposes and is further supported by the PF-127 hydrogel matrix.

### 4.5 Practical recommendations for adopting this method

For laboratories considering this approach, we offer several practical recommendations. First, restrict critical recordings to the first 30 minutes when motion reduction is most robust. Second, always include hydrogel-only controls to enable within-experiment assessment of any anesthetic effects on neural dynamics. Third, use the dual-flag QC system (`MOVEMENT_VALID` and `ROI_ELIGIBLE`) to ensure transparent reporting of which recordings were included in each analysis and why. Fourth, report motion metrics alongside calcium imaging results, since motion artifacts represent a potential confound even in well-immobilized preparations.

The entire analysis pipeline, including motion tracking, quality control, statistical comparison, calcium imaging processing, and ROI refinement, is implemented in self-contained MATLAB scripts with no external dependencies beyond the Statistics and Machine Learning Toolbox, Image Processing Toolbox, NoRMCorre, and Bio-Formats. All analyses use fixed random seeds for reproducibility. Code is available at https://github.com/arisakadrosophilalab/Larval-Immobilization-Analysis.

### 4.6 Limitations

Several limitations should be noted. Sample sizes are modest (*N* = 15 per group for movement; *N* = 23 control and *N* = 21 experimental for calcium), and all experiments used second-instar larvae. The optimal ether exposure parameters (duration, concentration, temperature) were not systematically varied; the 5-minute protocol used here was selected based on published protocols and may not represent the optimal trade-off between immobilization and neural perturbation. The lower SNR observed in experimental calcium recordings warrants further investigation to determine whether this reflects a genuine effect of ether on calcium dynamics, reduced transient amplitude due to partial neural suppression, or an artifact of other preparation differences. All data were collected in a single laboratory using a single microscope configuration, and broader reproducibility remains to be established. Recovery trials did not track larvae beyond the 30-minute observation window, so survival rates and long-term viability after ether exposure were not formally assessed.

Importantly, this study did not include independent validation of neural activity fidelity. No electrophysiological recordings, stimulus-evoked responses, or behavioral correlations were used to confirm that the calcium transients detected under ether exposure correspond to physiologically meaningful neural activity. The comparison was limited to hydrogel-only versus ether + hydrogel; an ether-alone condition was not included, and no comparison to microfluidic or other gold-standard immobilization methods was performed. The hydrogel-only baseline was chosen as the practical accessible standard, but this means the results demonstrate improvement over an accessible method rather than superiority to the best available techniques.

Calcium imaging comparisons at the ROI level benefit from large ROI counts but should be interpreted with caution, as ROIs within a sample are not independent. Sample-level comparisons, which better respect the experimental unit, showed weaker effects that did not always survive correction for multiple comparisons. Mixed-effects models that formally account for the nested data structure (ROIs within samples) would provide more rigorous inference and should be considered in future analyses.

The three motion metrics used (mean speed, median step, jitter ratio) were selected to capture complementary aspects of residual motion, but their direct relationship to calcium imaging signal quality was not formally validated. Future work correlating motion metrics with downstream signal fidelity would strengthen their use as quality indicators. The neuropil correction coefficient (0.7) follows the convention established by Chen et al. [2013] and adopted in subsequent GCaMP studies including wide-field applications [Dana et al., 2019], but has not been independently calibrated for the specific optical configuration and tissue geometry of our preparation.

We also note that because ether is a volatile anesthetic that could affect baseline fluorescence through changes in intracellular calcium homeostasis or pH effects on GCaMP, global signal regression (which removes condition-wide fluorescence shifts) would obscure any global excitation or suppression of activity. Our analysis can detect relative differences between ROIs but not uniform changes across the entire field of view. Studies specifically interested in global activity levels should be designed with this limitation in mind.

### 4.7 Future directions

Future work will focus on systematic optimization of ether exposure parameters, including dose-response characterization across developmental stages. Longer recordings (*>* 60 minutes) would help characterize the full recovery dynamics and determine whether neural activity normalizes over time. Direct comparison of spontaneous activity patterns between matched ether-treated and untreated preparations, ideally using simultaneous electrophysiology, would provide a more complete understanding of anesthetic effects on circuit function. Application to additional genotypes, GAL4 drivers, and calcium indicators will determine the generalizability of these results. Finally, post-imaging behavioral and viability assessments would help quantify the physiological impact of the combined protocol.

## 5 Conclusions

A multimodal immobilization approach combining PF-127 hydrogel with brief diethyl ether exposure (5 minutes at 25°C) significantly reduced residual motion during wide-field imaging of second-instar *Drosophila* larvae compared to hydrogel alone. Over the full recording duration (∼60 minutes), all three core motion metrics showed significant reductions after FDR correction (*q <* 0.001), with effect sizes ranging from medium to large (|*g*| = 0.80–1.36). In an exploratory analysis of the first 30 minutes, effects were uniformly large (|*g*| = 1.10–1.51), consistent with stronger initial chemical immobilization that is maintained by mechanical restraint as anesthetic effects diminish.

The approach is practical and accessible, requiring only standard laboratory supplies without specialized microfabrication. A dual-flag QC system (MOVEMENT VALID, ROI ELIGIBLE) supports transparent reporting and principled gating of calcium analysis to reliable recordings. Calcium imaging in both conditions demonstrated successful ROI detection with biologically plausible event rates, though experimental preparations showed lower SNR with event rates that were not suppressed relative to controls. Together, these results establish a practical workflow and transparent analysis framework for larval calcium imaging, with quantitative evidence that brief ether pre-treatment meaningfully improves immobilization quality.

## Data Availability

All analysis code is available at https://github.com/arisakadrosophilalab/ Larval-Immobilization-Analysis. Primary scripts include: `analyze_motion_and_QC.m` (motion analysis, 1 Hz GFP data), `analyze_motion_and_QC neuro.m` (motion QC, 33.33 Hz calcium data), `compare_motion_groups.m` (statistical comparison; the first-30-minute exploratory analysis is a configuration option of this script), `COMPLETE_PIPELINE_RUN_ME.m` (calcium imaging pipeline), `Assess_ROI_Quality.m` (signal quality assessment), `ROI_REFINEMENT.m` (multi-criteria ROI refinement), and `COMPARE_NEURO_CONDITIONS.m` (calcium imaging group comparison). Software environment: MATLAB R2024b (MathWorks), NoRM-Corre (motion correction), Bio-Formats (file I/O), Statistics and Machine Learning Toolbox, Image Processing Toolbox. Raw data are available from the corresponding author upon reasonable request.

## Author Contributions

**David Reynolds:** Conceptualization, Methodology, Software, Validation, Formal analysis, Investigation, Data curation, Writing – original draft, Writing – review & editing, Visualization, Supervision, Project administration.

**Eliana Artenyan:** Methodology, Investigation, Resources, Writing – review & editing.

**Hayk Nazaryan:** Investigation, Resources, Writing – review & editing.

**Edward Shanakian:** Investigation, Resources, Data curation, Writing – review & editing.

**Elva Chen:** Investigation, Data curation, Writing – review & editing.

**Vaheh Abramian:** Investigation, Data curation, Writing – review & editing.

**Ava Ghashghaei:** Investigation, Visualization, Writing – review & editing.

**Kian Sahabi:** Investigation, Data curation, Writing – review & editing.

**Fuad Safieh:** Investigation, Data curation, Writing – review & editing.

**Nicolas Momjian:** Investigation, Validation, Data curation, Writing – review & editing.

**John Sunthorncharoenwong:** Investigation, Data curation, Writing – review & editing.

**Javier Carmona:** Resources.

**Katsushi Arisaka:** Resources, Supervision, Writing – review & editing.

## Acknowledgments

We thank Alex Petrossian, Ariane Cazaubon, Calista Miramontes, Peter Elias, Tyler Humphries, and George Naddour for technical assistance and helpful discussions. We thank the Mark Frye laboratory (UCLA) for providing the R57C10-GAL4 and UAS-jGCaMP8f fly lines, and the David Krantz laboratory (UCLA) for providing the UAS-mCD8-GFP line.

## AI Disclosure

The authors used Claude (Anthropic) for assistance with manuscript preparation, editing, and reference verification. All content was reviewed and verified by the authors, who take full responsibility for the accuracy and integrity of the work.

